# Gut microbiome-derived prolyl peptidases from *Segatella copri* and *Stenotrophomonas maltophilia* degrade immunogenic gliadin peptides and restore intestinal barrier integrity in a Celiac disease model

**DOI:** 10.1101/2025.10.03.680265

**Authors:** Bhagyashree Karmarkar, Dhiraj Dhotre

## Abstract

Celiac disease (CeD) is an autoimmune enteropathy triggered by gluten-derived peptides that resist gastrointestinal digestion, notably the proline-rich 33-mer and 11-mer gliadin epitopes. Here, we describe a rational, data-centric strategy to identify gut microbiome-derived prolyl peptidases capable of degrading these immunogenic peptides. Integrating metagenomic mining with structure-based in silico screening, we identified two novel enzymes PSP692 and PSP464 from *Segatella copri* and *Stenotrophomonas maltophilia*, respectively. Recombinant expression and enzymatic characterization confirmed their activity under physiologically relevant conditions: PSP692 efficiently degrades the 33-mer at pH 6, while PSP464 targets the 11-mer at pH 4. Functional assays using CaCo-2 monolayers demonstrated that enzymatic degradation of gliadin epitopes significantly restored expression of tight junction proteins (ZO-1 and Occludin), reduced IL-6 secretion, and improved barrier integrity. These findings establish a foundational strategy for the discovery of microbiome-derived glutenases and provide a compelling case for combinatorial enzyme therapy to mitigate gluten immunotoxicity in CeD and related disorders.

## Introduction

Celiac disease (CeD) is a chronic autoimmune disorder triggered by the ingestion of gluten, a storage protein found in wheat, barley, and rye^1^. In genetically predisposed individuals^2–4^, gluten-derived peptides which are rich in proline and glutamine residues^5,6^, evade complete proteolysis in the gastrointestinal tract, leading to the activation of the adaptive immune system and subsequent duodenal inflammation. The only currently accepted treatment is a strict lifelong gluten-free diet (GFD), which is difficult to maintain and often fails to prevent inadvertent gluten exposure. Consequently, there is a pressing need for adjunctive therapeutic strategies that can mitigate gluten toxicity in situ, particularly by degrading immunogenic gluten peptides prior to their interaction with the mucosal immune system.

The human gut microbiome plays a pivotal role in both the exacerbation and alleviation of CeD pathogenesis^7^. Dysbiosis, characterized by the expansion of pro-inflammatory taxa and loss of beneficial commensals, can potentiate disease activity by enhancing intestinal permeability and generating immunogenic gluten-derived peptides through unregulated proteolysis^8^. Conversely, commensal bacteria such as *Bacteroides*, *Prevotella*, and *Lactobacillus* spp. may harbor intrinsic prolyl-specific peptidases that contribute to gluten detoxification by cleaving proline-and glutamine-rich motifs into less immunogenic fragments^9^. Several studies now support that the functional potential of the microbiome rather than its taxonomic composition alone critically shapes host responses to dietary gluten and influences disease outcomes^10^. Leveraging this microbial enzymatic repertoire through metagenomic mining and structure-guided discovery has already uncovered novel peptidases. Recent synthetic biology approaches extend this concept further, demonstrating that genetically engineered commensal gut bacteria can be tailored to stably express certain enzymes in situ^11^, thereby providing sustained protection against inadvertent gluten exposure while other approaches use recombinant enzymes. Together, these insights highlight the gut microbiome as both a key contributor to CeD pathogenesis and a promising reservoir of therapeutic enzymes.

Several microbial enzymes, particularly prolyl endopeptidases (PEPs), dipeptidyl peptidase IV (DPPIV), and other serine proteases^12–14^, have been investigated for their ability to cleave the immunodominant 33-mer gliadin peptide (LQLQPFPQPQLPYPQPQLPYPQPQLPYPQPQPF). However, most of current research remains limited to the characterization of enzymes discovered through labor-intensive functional screening, which inherently constrains the pool of candidates advancing toward clinical application. In contrast, our study advocates for a more rational and data centric approach by integrating metagenomic data with in silico predictive tools to pre-select promising glutenases based on structural and functional criteria.

In this study, we leveraged metagenomic data and peptidase databases to identify and characterize microbial proteases with gluten-degrading potential. Candidate enzymes (PSP692 and PSP464) were selected based on structural similarity to known glutenases and their activity against gliadin epitopes as predicted from docking and molecular dynamics simulation. These two promising enzyme candidates were cloned and expressed recombinantly, followed by purification and biochemical characterization. Enzymatic assays confirmed their activity against synthetic gliadin peptides. Furthermore, immunological assays using Caco-2 cell line demonstrated a reduction in gliadin-induced inflammatory markers, indicating that the enzymatic cleavage products were rendered non-immunogenic. Notably, we evaluated enzyme activity against both full-length and truncated gliadin immunogenic peptides, including the 33-mer and 11-mer epitopes^15^. Our findings demonstrate that PSP692 exhibits robust activity against the full-length 33-mer, while PSP464 preferentially degrades the 11-mer fragment (GPQQSFPEQEA), highlighting their complementary substrate specificities. This dual-targeting strategy represents a departure from conventional single-enzyme approaches that target only the 33-mer and supports the development of enzyme cocktails tailored for comprehensive gluten detoxification.

Taken together, our study has validated two enzyme candidates derived from metagenomic data and peptidase databases which expands the repertoire of microbial glutenases and demonstrates the utility of our data centric approach over the conventional activity screens. These findings provide a foundational step toward the rational selection and development of complementary enzyme supplements that can degrade dietary gluten in vivo, thereby reducing the immunogenic burden in CeD patients.

### Data pre-processing

Data submitted under NCBI BioProject ID PRJNA757365^16,17^ and PRJNA486782^18^ were downloaded and analysed with in-house methodology for taxonomic abundance profiling, quality assessment, assembly and binning, gene classification and translated protein search. Briefly read files of studies on CeD stool metagenomes of which datasets PRJNA757365 and PRJNA486782 are the only ones publicly available, were downloaded. The quality was assessed with FastQC (http://www.bioinformatics.babraham.ac.uk/projects/fastqc/) and it was determined that no read trimming is required. Taxonomic abundance was determined using MetaPhlAn2^19^, assembly was performed with MEGAHIT^20^, binning was done using MaxBin2^21^. Bins identified as belonging to organisms in higher abundance in non-CeD individuals by CheckM^22^ were submitted to Kraken2^23^ for gene classification. Contigs containing coding regions were identified in the ‘classified’ output of Kraken2 and presence of ORFs was determined using the eggnog-mapper module of HumanNv3.6^24^. The RAPSearch2^25^ module was then utilized for translated protein similarity search using the UniRef90 database. Proteins from the output fitting our in-house criteria (Figure 1) were selected for in silico evaluation.

**Figure 1:**
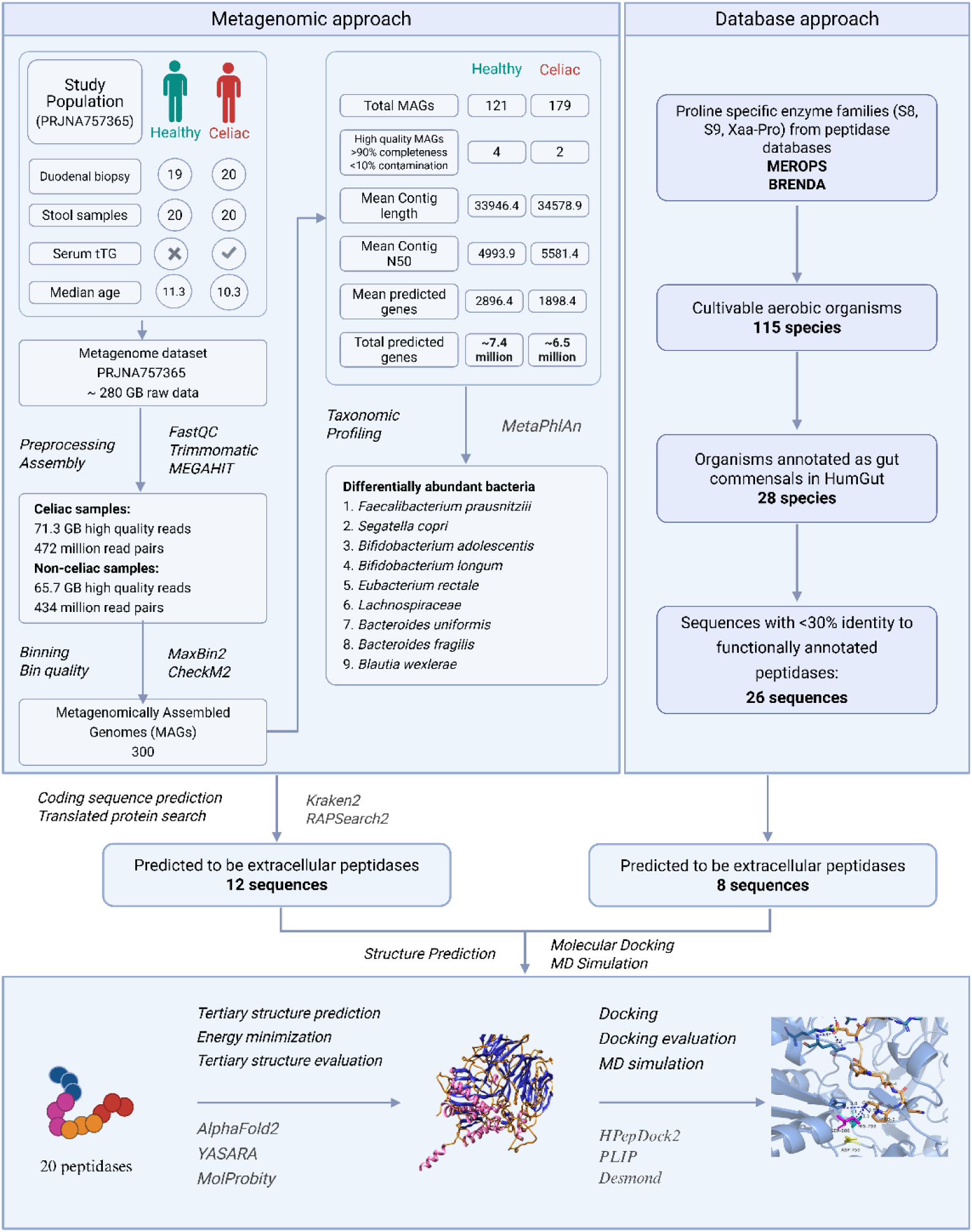
Reads of Celiac metagenome dataset PRJNA757365^16,17^ were checked for quality using FastQC and assembly was done using MEGAHIT^20^. The contigs were sorted into bins using MaxBin2^21^ which identified 121 bins i.e organisms present in healthy control samples and 179 such bins in samples from Celiac patients. Contigs likely containing coding sequences were identified by Kraken2^23^ and the presence of an open-reading frame (ORF) was determined by the eggnog-mapper module of HumanN3.6^24^(supplementary file ST1). RAPSearch2^25^ was used to perform a translated alignment and from the result file, sequences annotated to belong to proline-specific peptidase enzyme families and are present in one of the differentially abundant organisms were selected. PSPs predicted to be extracellular, and thus more likely to have a role in degradation of gut luminal gluten/gliadin, by a consensus of three tools: SignalP6^56^, PrediSi^57^ and Phobius^58^ were submitted for tertiary structure prediction (ST2C). From the MEROPS^27,43^ and BRENDA databases, 26 PSPs present in cultivable organisms and annotated to be non-pathogenic gut commensals were used for in-silico analysis. Twelve peptidase sequences from metagenome and 8 from databases underwent in-silico screening to identify high potential candidates for experimental validation.

### Database analysis

To identify candidate gluten-and gliadin-degrading enzymes with no prior annotation for such activity, we analysed data from two databases: BRENDA^26^ (The Comprehensive Enzyme Information System) (https://www.brenda-enzymes.org/) and MEROPS (The Peptidase Database)(https://www.ebi.ac.uk/merops/)^27^. BRENDA was used to screen for hydrolases, specifically peptidases, based on their EC numbers, substrate specificity, and reported kinetic parameters. Enzymes with biochemical properties suggestive of activity against proline-and glutamine-rich substrates were shortlisted. In parallel, MEROPS was used to analyze peptidases at the family and clan level, with a focus on serine peptidases previously implicated in gliadin cleavage. MEROPS data on active site residues, substrate cleavage sites, and known inhibitors were used to further refine enzyme selection.

For all shortlisted enzymes, the source organism was identified and cross-checked against literature and publicly available microbiome datasets to confirm whether it is a cultivable, non-pathogenic commensal of the human gut. Only enzymes meeting these criteria were retained for downstream structural, in silico analysis, and experimental validation. This multi-tiered selection strategy enabled the identification of functionally probable, previously uncharacterized gluten-degrading enzymes from gut-relevant microbial sources.

### In-silico evaluation

Sequences of peptidases obtained from databases and metagenome data analysis were submitted to AlphaFold2^28^ for tertiary structure prediction. All the models were subject to energy minimization using SwissPDBviewer^29^ and evaluated using MolProbity^30^. The top ranked model of each peptidase was used for docking. Docking of predicted tertiary structure of peptidase was done using the HPepDock^31^ server. The ligands were selected from PDB, with the selection criteria being: peptide ligand is bound to either MHC or TCR and structure is determined by X-ray crystallography with a resolution < 2Å. The docked structures were evaluated using PLIP^32^ (Protein Ligand Interaction Profiler) and peptidases whose catalytic residues formed a donor hydrogen bond of < 3 Å with ligand were selected for MD simulation. Prediction of active site residues was done using GASS^33^ (http://gass.unifei.edu.br). After tertiary structure prediction and docking, structures wherein the enzyme catalytic residues were predicted to form a donor hydrogen bond of length less than 3Å were selected for MD simulation. MD simulation was performed with Desmond^34^ academic version 2021-1 via Schrodinger Maestro GUI.

### Cloning of prolyl peptidase genes

Genes encoding for PSP464 and PSP692 were amplified from the genomic DNA of *Stenotrophomonas maltophilia* MCC2084 and *Prevotella copri* DSM18205 respectively. Primer sequences used for amplification were as follow: i) PSP464-F 5’-ATTGGATCCATGCTCATCCGTCGAAC-3’ ii) PSP464-R 5’-CTAAAGCTTCTTGGCGTTCTTCACGTAG-3’ iii) PSP692-F 5’-CGGGATCCGATGAAAAGATTTCCAGTTTTTATTGC-3’ and iv) PSP692-R 5’-CCCAAGCTTCTTCAGATTCTGCTTGAACCAGTTA-3’. The amplified genes were cloned into a pET22b plasmid (Novagen Healthcare Pvt. Ltd.).

### Expression and purification of prolyl peptidases

Recombinant expression plasmids were introduced via transformation into *Escherichia coli* BL21(DE3) cells. Transformants were grown at 37°C, and induced in the presence of 0.5/1 mM IPTG at 18 °C overnight. Lysis was performed by pelleting down induced cultures and resuspending in SDS (final concentration 1% w/v) solution, followed by centrifugation to collect supernatant. Equal amount of protein from uninduced and induced cell lysate was analysed on a 12% SDS-PAGE gel, transferred to nitrocellulose membrane and probed with anti-His antibody (Cat no:2366T, CST Inc.) to confirm expression of full-length recombinant protein.

All purification steps were performed at 4 °C unless noted otherwise. Recombinant bacterial culture was pelleted down at 4000-5000g, supernatant was discarded and cells were lysed by resuspension in solution with final concentration of SDS at 1% w/v. The lysate was centrifuged at ∼12,000g for 30min and the supernatant was collected for purification. Volume of Ni-NTA magnetic beads (GenScript) added to suspension was one-third total volume of lysate, the mixture was then spun on rotospin at 15rpm for atleast 1 h. Elution was done using reaction buffers of recombinant enzymes, which did not contain any imidazole.

### Activity assay

Post-proline cleavage activity of the purified enzymes was assessed using two chromogenic substrates: Z-Gly-Pro-pNA (Sigma-Aldrich Chemicals Pvt. Ltd.) and a synthetic peptide substrate Z-PPF-pNA, which was custom-synthesized at >95% HPLC purity (Genetoprotein Pvt. Ltd.). Z-Gly-Pro-pNA was prepared as a 50 mM stock solution in dimethyl sulfoxide (DMSO) and stored at –20°C until use. Z-Pro-Pro-Phe-pNA was prepared as a 20mM stock solution in nuclease free water and stored at-20°C until use. Hydrolytic activity against Z-PPF-pNA was evaluated for enzyme PSP464 in 100 mM glycine-HCl buffer at pH 4.0, while Z-Gly-Pro-pNA hydrolysis by enzyme PSP692 was measured in 1X phosphate-buffered saline (PBS) at pH 6.0. Enzymatic cleavage of the pNA-linked substrates was monitored spectrophotometrically by measuring the increase in absorbance at 410 nm, which corresponds to the release of p-nitroaniline upon substrate hydrolysis.

Each reaction was carried out in a final volume of 100 µL in thin-walled 0.5mL PCR tubes, and contained a final enzyme concentration of 1 µM. Substrate and enzyme solutions were added such that their combined volume did not exceed 50% of the total reaction volume. The concentration of DMSO was carefully maintained below 0.5% in all assays to minimize solvent effects on enzyme activity. Absorbance values obtained were used to calculate enzymatic activity based on a standard curve generated with known concentrations of para-nitroaniline in the respective assay buffers. Background absorbance from substrate auto-hydrolysis was accounted for by subtracting the optical density (OD410) of control wells containing substrate without enzyme. All measurements were performed in triplicate to ensure reproducibility.

### Cell culture maintenance and assay method

CaCo-2 cell line was obtained from repository of National Centre for Cell Science, Pune. Complete culture medium was composed of DMEM medium supplemented with 10% FBS and 1% non-essential amino acids, incubated in humidified chamber at 37°C and 5% CO_2_. All media components were purchased from Gibco, ThermoFisher Scientific, Inc. Cultures were split in a 1:3 ratio once every 48-72h i.e. when ∼80-90% confluence was achieved. For assays involving measurement of cytokine secretion and gene expression levels, cells were seeded at an initial density of 5*10^4 cells/well of a 24-well culture plate and maintained in 1mL complete medium for atleast 72h prior to assay. For monolayer permeability assay, cells were grown on membrane for a minimum of 28 days until transepithelial resistance of each well was a minimum of 500 Ω/cm^2^. During this 28 day culture period medium was replaced once every two-three days. On the day of assay, cell culture medium was aspirated and replaced with: culture medium (control), 1mg/mL of undigested 33-mer (LQLQPFPQPQLPYPQPQLPYPQPQLPYPQPQPF) or 11-mer (GPQQSFPEQEA), 1mg/mL of peptides digested with PSP692 or PSP464 in a 1:500 molar ratio. Synthetic peptides were purchased from Genetoprotein Pvt. Ltd. at an purity >95% as verified by HPLC.

### Measurement of IL-6 secretion

Previously collected culture supernatant were thawed (not more than one freeze-thaw cycle) and used for IL-6 estimation in pg/mL by ELISA as per manufacturers instruction (Cat. No: 900-M16 Human IL-6 Mini ABTS ELISA Development Kit, Mfg: PeproTech, Inc.). Readings were recorded at 405nm and 650nm at 10 min intervals for total time of 40 minutes. Time point where readings of assay standard displayed maximum linear corelation between concentration and absorbance was used.

### Quantitation of ZO-1 and Occludin transcript levels

The effect of 33-mer and 33-mer digested by PSP464 and PSP692 on ZO-1 and OCLN mRNA expression was investigated by real-time PCR (qRT-PCR) based on the SYBR Green method. Primer sequences were designed in-house and are listed in **Table 1**, primers were purchased from IDT, Inc. GAPDH was utilized as endogenous control. Cells were exposed to either 1mg/mL 33-mer gliadin/11-mer (PDB ID: 5KSB_I) epitope or to peptides of same concentration pre-digested with purified enzymes, either PSP464 or PSP692. Incubation period of assay was 4h, at the end of which cells were harvested with 0.25% trypsin-EDTA, washed with sterile 1X PBS and resuspended in equal volumes of Trizol (ThermoFisher Scientific Inc.) for RNA extraction per manufacturers protocol. The quality and quantity of total RNA were detected using a NanoDrop® spectrophotometer. Equimolar RNA was used to synthesise cDNA using Verso cDNA kit (ThermoFisher Scientific Inc.). SYBR Green Master mix (ThermoFisher Scientific Inc) alongwith cDNA and primers were used to perform qPCR, 2^-ΔΔCt^ method was used to interpret the results.

**Table 1:**
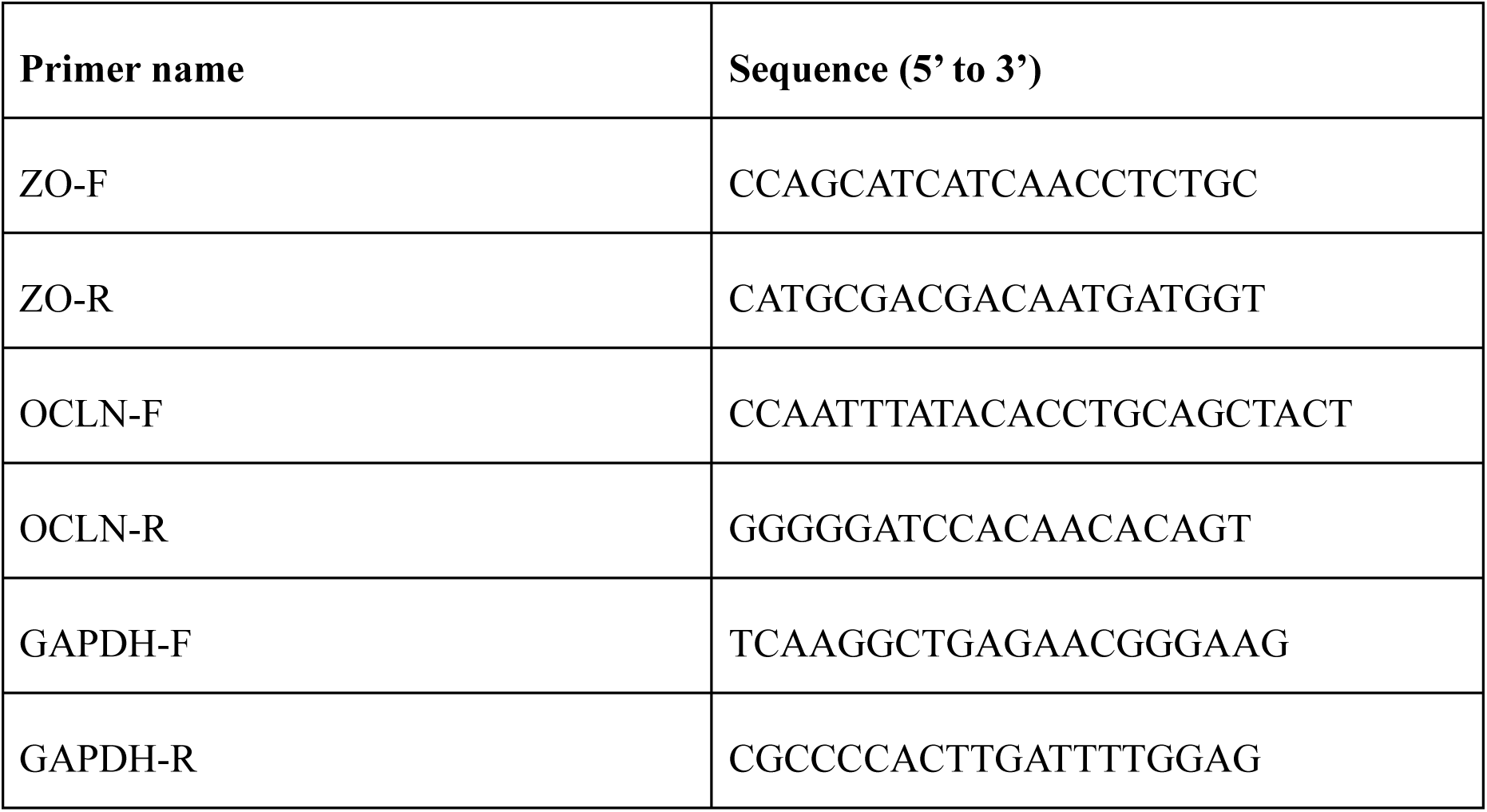
Sequences of primers designed in-house for RT-qPCR.

### Measurement of monolayer permeability

Transepithelial electric resistance (TEER) was used to monitor the integrity of the epithelial monolayer and was determined using a Millicell ERS-1 volt-ohm meter according to the manufacturer’s instructions. TEER values were calculated as ohms × cm^2^. Monolayers reaching TEER values >500 Ω · cm^2^ were considered to have an appropriate barrier function and were used for further study. Paracellular permeability was measured as Dextran-FITC 4kDa (SRL Pvt. Ltd.) fluorescence intensity on apical side.

### Quantitation of ZO-1 expression at CaCo-2 cell surface

Changes in expression of ZO-1 at cell surface were determined by confocal microscopy. Cells were incubated for 4h with 33-mer only or enzyme digested 33-mer, cells for microscopy were grown in 12-well removable chamber slide (Ibidi Pvt. Ltd.). At the end of incubation period cells were washed with 1X PBS and fixed in 4% para-formaldehyde in 1X PBS for 10-15 min. This was followed by washing with 1X PBS, following which blocking was done for 1h at room temperature in 5% BSA. Then 100 µl of a 1:2500 dilution of anti-ZO1-AlexaFluor647 (ThermoFisher Scientific Inc.) in 3% BSA in 1X PBS was added to each sample well and incubated overnight at 4℃ in dark. Antibody solution was aspirated, slides were washed with 1X PBS; and a 1:10,000 dilution of DAPI was added to each well 10-15 min prior to washing followed immediately by addition of 90% glycerol as mountant. Cells stained with DAPI only were used as negative control.

### Statistics

Data were analyzed with GraphPadv8.3(San Diego, CA) software. One way ANOVA test and Tukey’s post-test were used to determine statistical significance. A p-value of <0.05 was significant. Data are expressed as means ± standard deviation (SD) of triplicate samples for all assays performed.

## 3. Results

### 3.1 Bioinformatic analyses and in-silico screening

From the bioinformatic analysis of metagenome dataset PRJNA757365 (**Figure 1**), 9 bacterial species were found to be differentially abundant between Celiac and non-Celiac samples viz. *Faecalibacterium prausnitzii*, *Segatella copri*, *Eubacterium rectale*, *Bacteroides uniformis*, *Bacteroides fragilis*, *Bifidobacterium adolescentis*, *Bifidobacterium longum*, *Blautia wexlerae*, and Lachnospiraceae bacterium (with a normalized relative abundance >1) (supplementary ST2A, ST2B). Over 200 Proline-specific peptidases (PSPs) from these nine bacterial species and from peptidase databases that fit our inclusion/exclusion criteria, underwent in-silico screening.

After application of selection criteria to candidates from peptidase databases and metagenome data analysis, cumulatively 20 protein sequences were predicted by InterPro^35^ to contain PSP domains. The rank 1 models from AlphaFold2^28^ tertiary structure prediction for all 20 protein sequences were docked with all 7 gliadin epitope ligands using the HPepDock^31^ and HDock^36^ server in site-specific docking mode. Based on the following i) formation of a donor hydrogen bond of length <3A by catalytic serine with a peptide bond of the ligand (ST2F), ii) MM/GBSA of the docked structure (ST2D) and iii) ligand RMSD, protein-ligand contacts and ligand-protein contacts and percent occupancy of catalytic residues determined from MD simulation (ST2E); PSP464 and PSP692 sequences were identified as promising candidates for experimental validation.

Phylogenetic tree was constructed in MEGA12^37^ to compare our recombinant peptidases with bacterial peptidases that are validated to have gluten degrading activity (**Figure 1A**) and with various peptidases predicted from metagenomic data and annotated in MEROPS(**Figure 1B**). The bootstrap values were calculated from 500 replicates. All docking results were visualized in UCSF Chimera1.19^38^. MM/GBSA was determined using HawkDock server^39^. PDB2PQR and APBS functions were done using Chimera plug-ins. APBS result files were visualized in PyMol. To enable direct structural comparison, all molecular visualizations generated in PyMOL were aligned and displayed using identical view tuples (ST2G), ensuring uniformity in the spatial presentation of active sites and binding regions. Similar data analysis was applied to PRJNA486782, however none of the metagenomes obtained were of sufficient completeness as to be suitable for further screening.

We found that the gene encoding PSP692 from metagenome reconstructed genome of *S. copri* has identity of 80-85% to reference genome of *S. copri* DSM18205. The gene coding for PSP464 determined from the whole genome submission for *S. maltophilia* NCTC10257 (GenBank: GCA_900186865.1) is >90% identical to a gene from *S. maltophilia* WND-1. We further determined from phylogenetic analysis of protein sequences of PSP464 and PSP692 that our query peptidases cluster separately from previously validated gluten degrading enzymes. We further determined from phylogenetic analysis of protein sequences of PSP464 and PSP692 that our query peptidases cluster separately from previously validated gluten degrading enzymes. Sequences used for construction of phylograms are present in supplementary information. The clade formation observed in **Figure 2B** indicates that PSP sequences group according to enzyme class rather than source organism. This is exemplified by PSP692, derived from *S. copri*, which clusters with the DPPIV from *B. thetaiotaomicron* rather than with other peptidases from *Segatella*. Thus while all proline-specific peptidases roughly belong to S8/S9 class of enzymes their sub-class determines their substrate specificity which also underlies the ability of enzyme to abrogate immunogenicity of gliadin.

**Figure 2:**
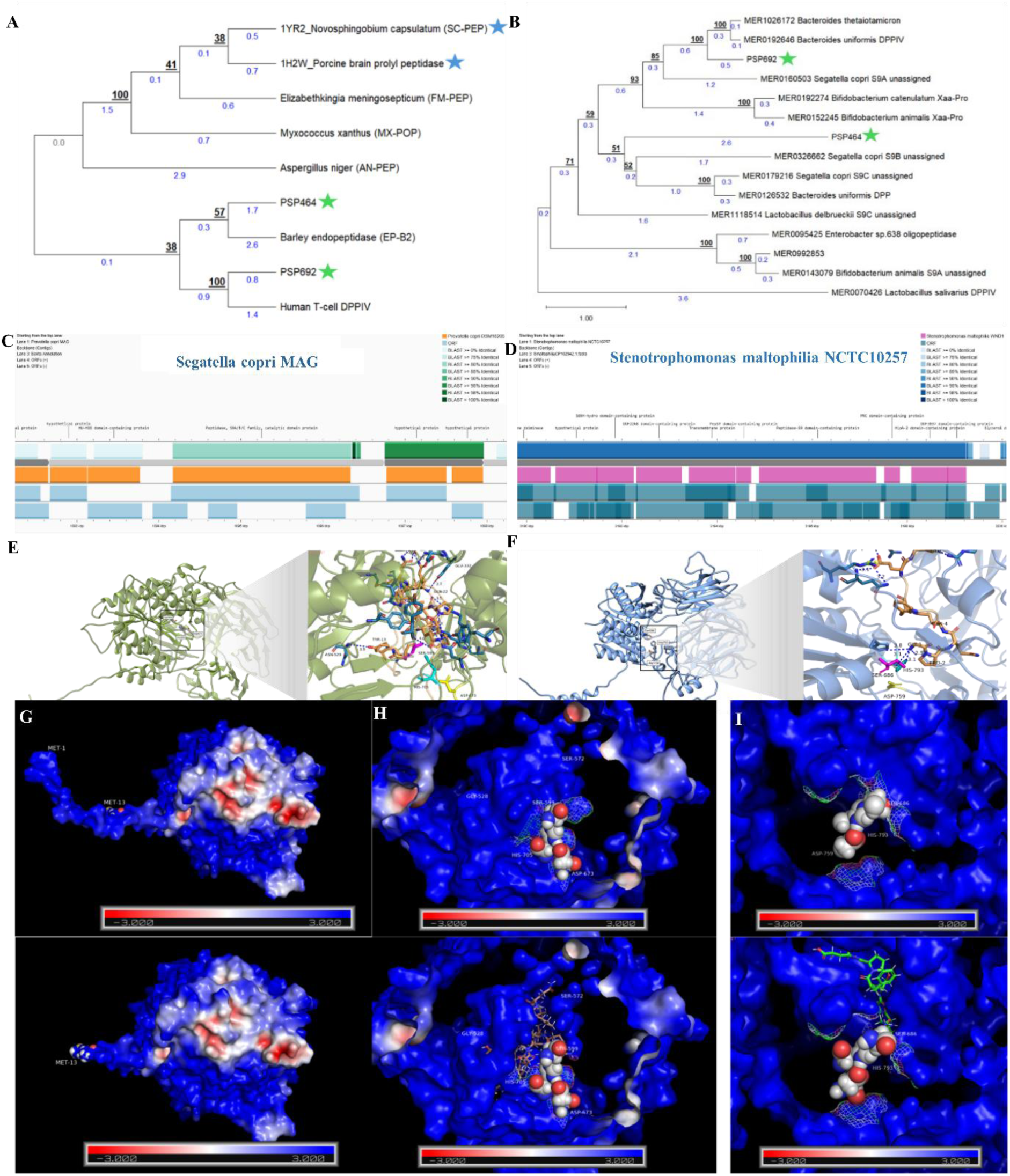
A) A phylogenetic tree constructed from query sequences(green star) of PSP464, PSP692 along with experimentally validated gluten-degrading enzymes (GDEs), GDEs with PDB structure (blue star), and metagenome derived PSPs that are annotated in MEROPS but not validated. For metagenome derived PSPs, bacteria from the top 10 genera of core human gut microbiome in UHGG^59^ annotated to be part of healthy human gut per HumGut^60^ were considered. **B)** De-duplicated and merged bins annotated as *Segatella copri* by CheckM^22^ was used to construct a genomic map annotated by the Bakta module of Proksee^61^. Location for PSP692 in reconstructed genome was determined by BLAST of contigs annotated by Bakta^62^ to be S9/S8/Xaa-Pro peptidase. **C)** The genome of *Stenotrophomonas matlophilia* NCTC10257 was downloaded from NCBI and used to construct genome map in Proksee^61^, this was then aligned to genomic standard strain *S. maltophilia* WND-1. **D)** Docking of PSP692 with 33-mer gliadin and a zoomed in view show the predicted active site: Ser599 (magenta), Asp673 (yellow) and His705 (cyan). Hydrogen bond interactions for docking of PSP692 with each of tested ligands alongwith length of the H-bond and binding energy are tabulated. **E)** Docking of PSP464 with 11-mer gliadin and a zoomed in view show the predicted active site: Ser686 (magenta), Asp759 (yellow) and His793 (cyan). Hydrogen bond interactions for docking of PSP464 with each of tested ligands alongwith length of the H-bond and binding energy are tabulated. **F)** Putative sites of ligand cleavage upon docking with PSP464 or PSP692 based on bond torsional energies determined from molecular dynamics simulation in Desmond. **G)** Top: Exterior view of APBS of only PSP692 at pH 6. Bottom: Exterior view of APBS of PSP692-33-mer docked complex at pH6. **H)** Top: View of active site of only PSP692 at pH 6, surface color is based on electrostatic charge distribution. Bottom: APBS of active site of PSP692 docked with 33-mer gliadin at pH6. **I)** Top: View of active site of PSP464 enzyme at pH 4. Bottom: View of active site of PSP464 docked with 11-mer gliadin at pH 4. Results of PLIP evaluation of PSP692-33-mer and PSP464-11-mer docked structures is present in supplementary at ST2F. All PyMol visualisations are available as session files in supplementary information alongwith the view tuples for each set present in supplementary at ST2G.

### 3.2 Effect of PSP692 and PSP464 on gliadin induced cytokine secretion and loss of barrier integrity

#### Determination of enzyme kinetics

Upon purification of recombinant enzymes by Ni-NTA affinity method, the enzyme kinetics and post-proline cleavage ability were determined by using two synthetic substrates viz. Z-GP-pNA and Z-PPF-pNA at variable concentrations. Reactions for PSP464 were carried out at 37°C in pH 4 buffer of glycine-HCl while reactions for PSP692 were carried out in pH 6 PBS buffer at the same temperature. The kcat values for PSP692 and PSP464 against the chromogenic gluten motif analogue, as calculated from a velocity versus substrate profile over 200–1000 μM substrate, were found to be 56.5 s–1 and 7 s–1, respectively (**Figure 3B**). The kcat value for PSP464 indicate a low turnover rate while PSP692 demonstrates an average to above average catalytic rate based on values reported for other non-mutated prolyl oligopeptidases.

**Figure 3:**
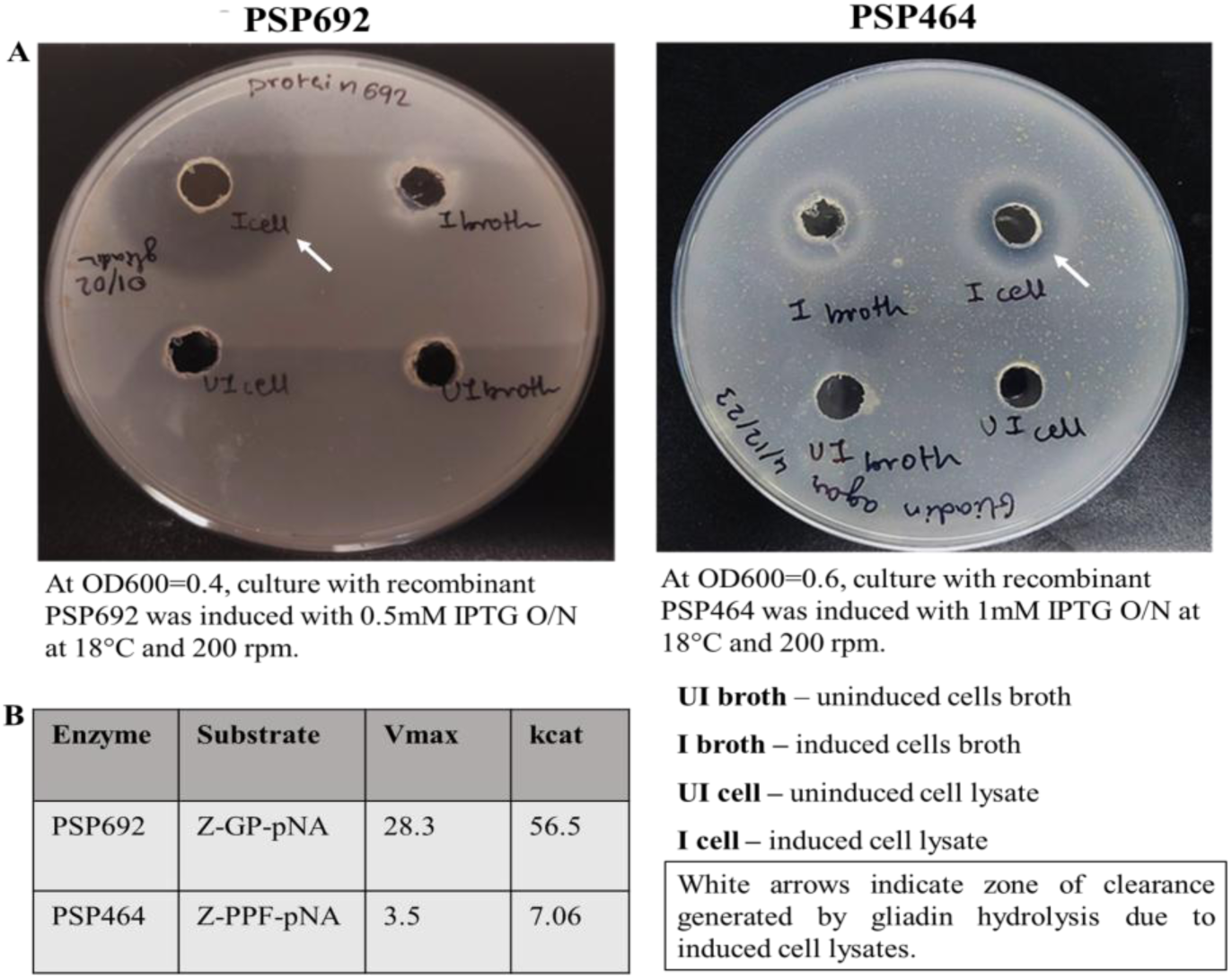
A) The retention of native gliadin degrading activity by recombinant PSP692 and PSP464 was determined qualitatively by plating 200ul of induced cell lysate in wells made in gliadin agar plate. The uninduced cell lysate was plated as control. The agar plates were incubated overnight at 37°C and as indicated by white arrows a zone of clearance shows that recombinant proteases are soluble and able to degrade gliadin. Some gliadin degrading activity is also observed in the broth of induced PSP464 culture indicating that it is secreted. **B)** The catalytic efficiency (kcat) for both PSP692 and PSP464 for synthetic peptides modified at C-terminus with p-nitroanilide were observed from a linear fit of velocity vs substrate concentration profile as no saturation was observed up to 1 mM substrate. The absorbance was quantified at 405nm as described in Methods. Rate of product formation was determined with a standard curve of p-nitroanilide in respective reaction buffers of PSP692 and PSP464.

#### Expression of ZO-1 and Occludin

The CaCo-2 cells upon reaching 80-90% confluence were exposed to either 1mg/mL of unidgested or enzyme digested peptide. After exposure to gliadin epitopes for 45 minutes the RNA was harvested and immediately used to synthesise cDNA. From the qPCR results we observed that PSP464 was able to significantly degrade the 11-mer epitope as evidenced by the restoration of ZO-1 transcript levels. Similarly, PSP692 was able to significantly degrade the 33-mer peptide, leading to restoration of the transcript levels of key barrier proteins, viz. ZO-1 and Occludin (**Figure 4A**). These findings suggest that PSP464 and PSP692 may play a role in ameliorating barrier function by reducing the immunogenic effects of gliadin-derived peptides.

**Figure 4:**
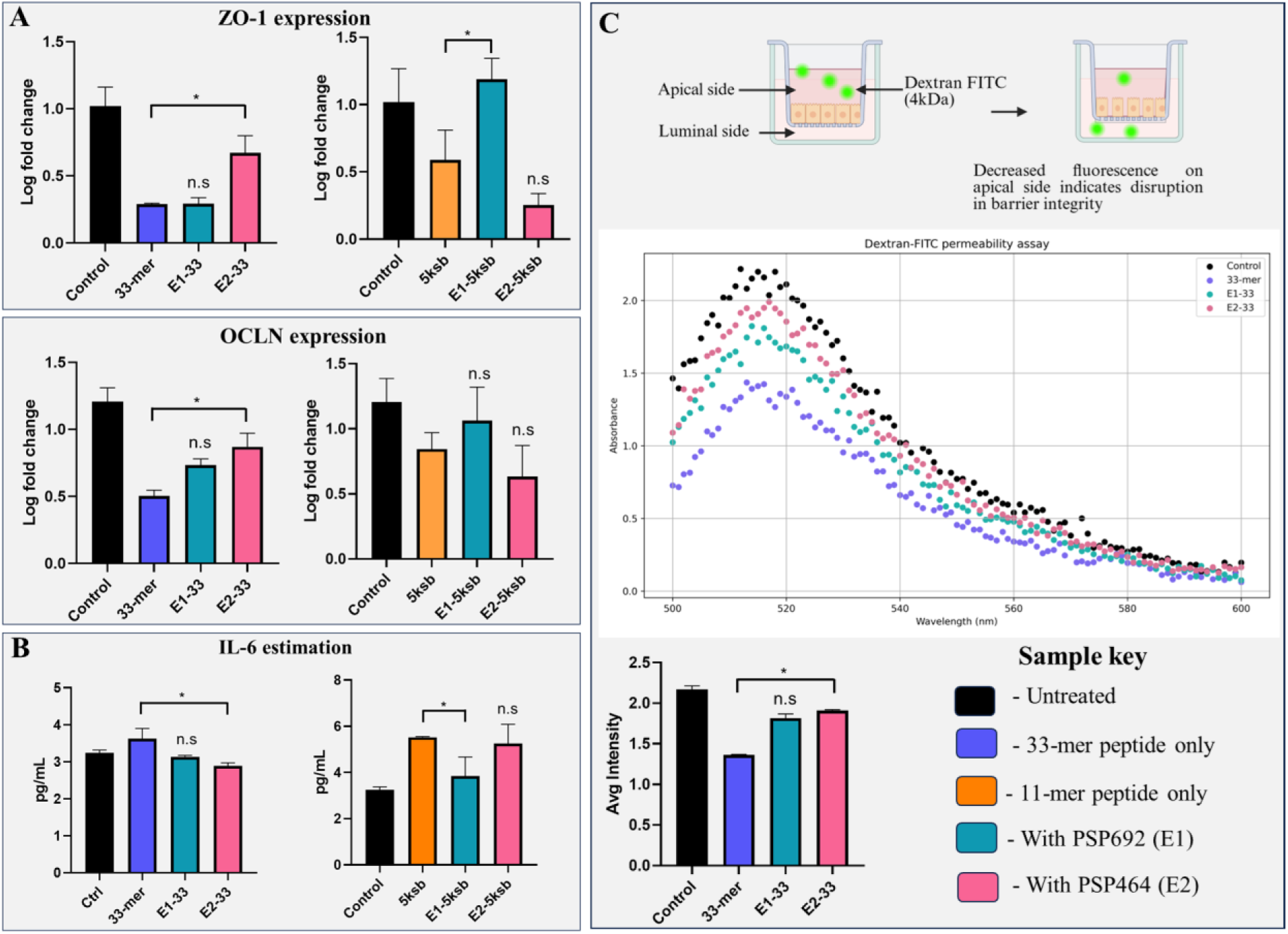
A &. **B)** Panels show the levels of ZO-1 and Occludin (OCLN) transcripts in cells incubated for 45 mins with undigested and enzyme-digested 33-mer and 11-mer gliadin epitopes. PSP692 is able to degrade 33-mer to an extent that significantly restores ZO-1 and OCLN expression while PSP464 is able to degrade 11-mer to an extent that significantly restores ZO-1 expression. **C)** Secretion of IL-6 into culture supernatant was determined by sandwich ELISA, as per manufacturer’s instructions the difference in absorbance at 650nm and 405nm was used to calculate IL-6 concentration by extrapolation from a standard curve. PSP692 degraded 33-mer while PSP464 degraded 11-mer gliadin to an extent that results in significant decrease in IL-6 secretion. **D)** Barrier function of CaCo-2 cells cultured on Transwell membrane for 28 days was determined as a function of 4kDa Dextran-FITC fluorescence in apical chamber, measurement range of 500-600nm. Disruption of barrier integrity leads to loss in signal intensity. Treatment of 33-mer with PSP692 prevents barrier disruption. For all assays, reactions were performed in triplicate, statistical significance (*= p <0.05) was determined using one-way ANOVA and Tukeys post-test. Error bars represent mean ± SD.

#### Inflammatory cytokine estimation

The secretion of IL-6 has been well-established as an important marker of gut inflammation based on Celiac patient duodenal biopsy derived organoids. At a 1:500 molar ratio of enzyme: peptide, PSP692 and PSP464 were able to degrade the 33-mer and 11-mer respectively (**Figure 4B**), to an extent that significantly reduces IL-6 secretion into the culture supernatant. These results provide evidence that the peptidases have a direct effect on mitigating the inflammatory response induced by gliadin and its derived peptides, thus potentially offering therapeutic benefits in celiac disease.

#### Restoration of monolayer barrier integrity

The effect of PSP464 and PSP692 in modulating barrier integrity was estimated based on flux of a small molecule viz. Dextran-FITC 4kDa from apical to basolateral region of Transwell. Fluorescence emission from apical side of Transwell membrane was determined spectrometrically in the range of 500-600nm at 1nm intervals (**Figure 4C**). However, readings in the range of 515-525nm, which correspond with emission maxima of FITC, were considered for statistic comparison. Relative to untreated cells, the addition of the 33-mer peptide to the apical side of the CaCo-2 monolayer resulted in a significant increase in monolayer permeability, as indicated by a decrease in the fluorescent signal of apical side solution. This demonstrates that the 33-mer peptide compromises the integrity of the epithelial barrier. However, when 33-mer treated with PSP692 was introduced, the permeability of the monolayer was significantly restored, as indicated by the recovery of the fluorescent signal. PSP464 also demonstrated significant efficacy.

#### Effect of enzyme treatment on surface expression of ZO-1

To further investigate the protective effect of PSP692 and PSP464 on Caco-2 cells, confocal microscopy was employed to visualize ZO-1 protein, a critical component of tight junctions (**Figure 5**). ZO-1 is a critical component of intestinal tight junctions, functioning as a platform for the attachment of occludin and claudins at intercellular site and for adherence of actin cytoskeleton on cytoplasmic side; overall this assembly plays a pivotal role in the regulation of barrier function. Nuclear staining with DAPI was also used to visualize any potential changes in nuclear morphology due to treatment with recombinant enzymes.

**Figure 5:**
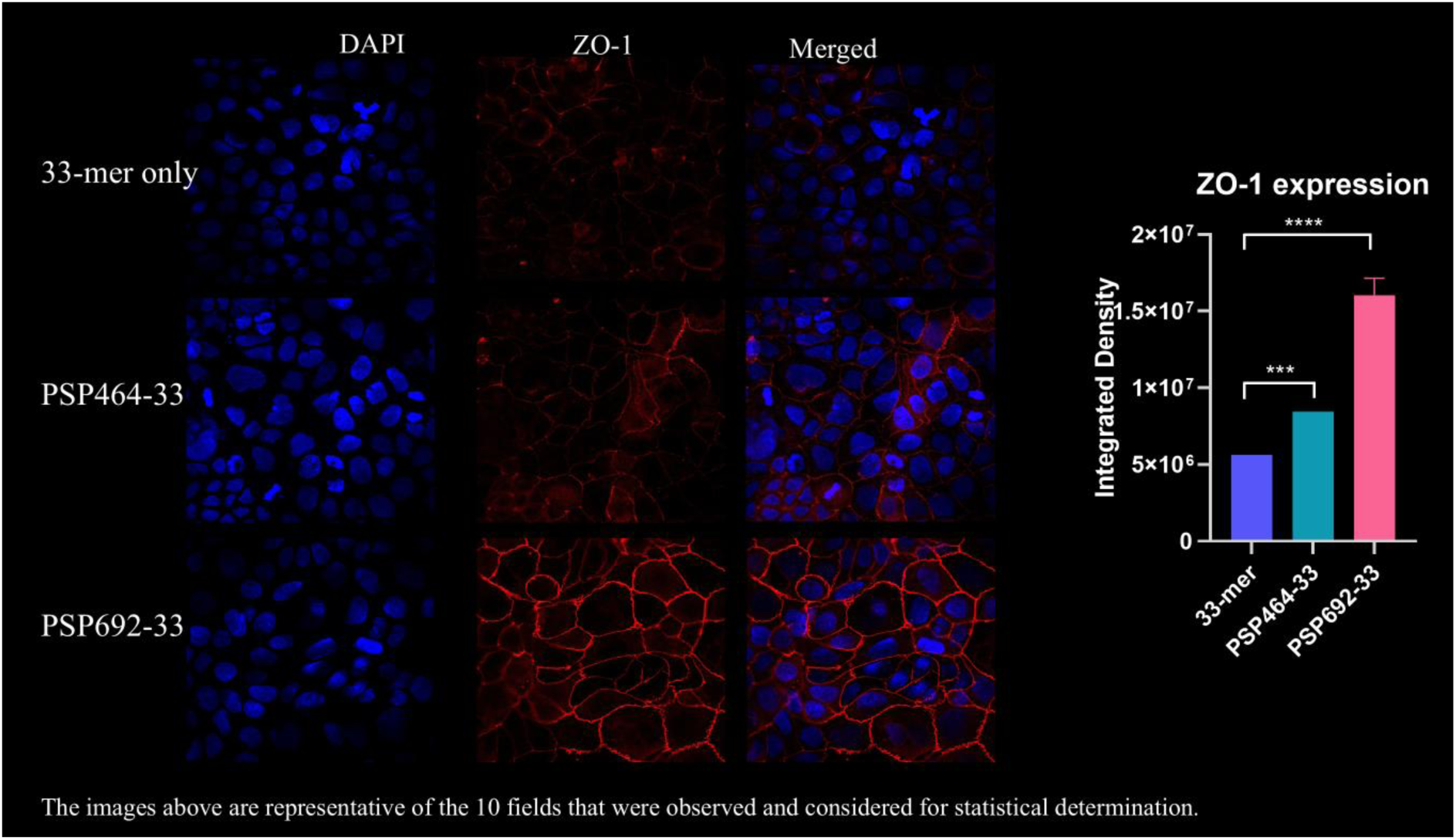
Quantitation of ZO-1 protein at the cell surface was determined by staining with a direct tagged anti-ZO-1 monoclonal antibody. For analysis of ZO-1 expression at cell surface in 33-mer undigested/pre-digested conditions ten images of each sample were acquired at random on Olympus FV3000 at 63X magnification. Images are representative of ten random fields acquired per sample.

The images for each sample were visualized in ImageJ using the Olympus viewer plugin. Fluorescence signal was determined for the full field. The averaged integrated density of all samples clearly demonstrates a very significant increase in expression of adhesion protein at tight junction sites when 33-mer gliadin is digested with either PSP464 or PSP692. This result demonstrates that the efficacy of PSP464 and PSP692 in restoring barrier function (by degrading 33-mer) is not restricted to increase in transcripts of ZO-1 but is also visible at the cell surface.

## 4. Discussion

Celiac disease is an autoimmune enteropathy characterized by chronic intestinal inflammation, initiated and sustained by immunogenic gluten-derived peptides, particularly the proline-rich 33-mer α-gliadin fragment that evade degradation by human gastrointestinal proteases. While host genetic susceptibility particularly HLA-DQ2/DQ8 plays a necessary role in disease onset^40^, not all carriers develop CeD evidenced by the absence of 100% concordance of disease development in monozygotic twins^41^, indicating the involvement of other environmental or microbial modulators^42^.

Traditional efforts in the search for glutenases have largely relied on labor-intensive functional screening of microbial isolates or environmental DNA libraries, which, while useful, are inherently time-consuming and inefficient. These approaches tend to yield a limited number of hits, often favoring enzymes that are abundant, easily expressed, or already well-characterized. In contrast, our approach leverages high-resolution metagenomic data from human cohorts viz. PRJNA757365 and PRJNA48678 (the only Celiac disease metagenome datasets that are publicly available), processed through a tiered pipeline optimized for both taxonomic and functional fidelity. Rather than relying solely on gene annotations or conserved domain matches, we employed metagenomic assembly (via MEGAHIT^20^) followed by taxonomic binning (MaxBin2^21^) and protein-level functional screening (via RAPSearch2^25^) using the UniRef90 database. This integrated pipeline enabled not only the identification of low-abundance but catalytically relevant genes, but also their linkage to specific taxa with high confidence. This was particularly advantageous in our study, where we were able to resolve strain-level genomes from *Segatella copri* and *Stenotrophomonas maltophilia*, and identify unannotated peptidases that shared structural similarity rather than sequence homology with known glutenases. Such structural convergence would have been missed by conventional alignment-based annotation strategies alone. Our method thus facilitated the discovery of functionally relevant yet previously unrecognized prolyl peptidases that are differentially abundant in non-CeD metagenomes, highlighting the power of structure-guided metagenomic mining for therapeutic enzyme discovery. We focused mainly on novel proline-specific peptidases (PSPs) (<30% similarity) enriched in metagenomic bins from non-CeD samples, hypothesizing that their relative abundance might correlate with protective phenotypes.

Phylogenetic analysis revealed that the two candidate enzymes identified in this study PSP692 from *Segatella copri* and PSP464 from *Stenotrophomonas maltophilia* cluster distinctly from previously characterized gluten-degrading enzymes which are validated experimentally (Figure 1B). Despite sharing conserved motifs with known proline-specific proteases, these enzymes do not cluster closely with functionally validated peptidases, highlighting their novelty and the untapped diversity of glutenases within the human gut microbiome. Both enzymes belong to the S9 (subtilisin-like serine proteases) and Xaa-Pro peptidase families, which are broadly implicated in the cleavage of proline-rich substrates but are rarely annotated with specific glutenase function in standard databases. For instance, the S9 family includes a wide spectrum of endopeptidases with overlapping catalytic triads (Ser-His-Asp), while Xaa-Pro dipeptidyl peptidases act at the N-terminal proline of peptides. However, the lack of detailed substrate-specific characterization for many members of these families often limits their therapeutic consideration.

To refine our selection, we integrated enzyme family classification from MEROPS^27,43^ and BRENDA with subcellular localization predictions, prioritizing extracellular enzymes likely to encounter dietary peptides in the gut lumen. AlphaFold2^28^-based structure prediction enabled resolution of high-confidence tertiary models for each enzyme, including the orientation of the catalytic triad and substrate-binding pocket. These structures were docked against a panel of immunogenic gliadin peptides namely the 33-mer (LQLQPFPQPQLPYPQPQLPYPQPQLPYPQPQPF) and shorter epitopes including 5KSB, 4OZI, and 5IJK using HPepDock^31^. Only those complexes exhibiting catalytically relevant interactions were selected for molecular dynamics (MD) simulations. These structural interaction criteria are well-established predictors of enzyme-substrate interaction^44,45^ and allowed us to prioritize enzyme-substrate pairs with high catalytic plausibility. The identification of PSP692 and PSP464, which had not been previously linked to gluten metabolism, underscores the power of this structure-guided metagenomic mining approach in uncovering novel, therapeutically relevant enzymes.

Following recombinant expression and purification, enzyme kinetics determined using Z-GP-pNA and Z-PPF-pNA respectively, revealed that PSP692 has a higher catalytic turnover (k_cat = 56.5 s⁻¹) compared to PSP464 (k_cat = 7 s⁻¹) of these particular substrates, however these substrates alone do not provide adequate insight into the ability to degrade native gliadin epitopes.

The functional consequences of enzymatic cleavage were evaluated in a CaCo-2 epithelial cell model, which mimics the intestinal barrier and responds to gliadin-induced epithelial stress^46,47^. In the untreated condition, exposure to the 33-mer (LQLQPFPQPQLPYPQPQLPYPQPQLPYPQPQPF) and 11-mer (GPQQSFPEQEA) led to pronounced epithelial disruption, characterized by a marked increase in IL-6 secretion, decreased mRNA expression of tight junction markers ZO-1 and Occludin, elevated monolayer permeability (via FITC-Dextran flux), and a loss of ZO-1 protein localization at intercellular junctions as visualized by confocal microscopy. These changes are in line with known immunopathological responses of intestinal epithelium to gluten-derived peptides in CeD.

Pre-digestion of the 33-mer with PSP692 significantly attenuated these adverse effects. IL-6 secretion was reduced to near-baseline levels (Figure 4C), and transcript levels of tight junction marker were restored significantly (Figure 4A-B). In comparison, PSP464-treated 11-mer also showed significant reduction in IL-6 levels and restoration of ZO-1, consistent with its action on a shorter epitope. Importantly, both enzymes restored transepithelial barrier integrity as measured by the decrease in FITC-Dextran permeability (Figure 4D), with PSP692 showing superior efficacy, consistent with its preference for the full-length 33-mer.

Confocal microscopy further confirmed these results at the protein level. In the untreated condition, ZO-1 staining was decreased, reflecting junctional disruption. In contrast, both PSP692-and PSP464-digested peptide treatments showed pronounced relocalization of ZO-1 to the cell membrane. Quantitative image analysis revealed a significantly higher fluorescence intensity in PSP692-treated cells compared to PSP464 (Figure 5), suggesting a more complete restoration of epithelial polarity and tight junction integrity. These findings collectively indicate that while both enzymes mitigate gliadin-induced epithelial injury, PSP692 demonstrates a more potent protective effect, likely due to its higher turnover and capacity to degrade the immunodominant 33-mer.

The two enzymes characterized in this study, PSP692 and PSP464, exhibit differential but complementary activity profiles. PSP692 demonstrated strong activity against the full-length 33-mer gliadin peptide, which is the immunodominant epitope. In contrast, PSP464 showed more efficient degradation of the shorter 11-mer gliadin fragment. This finding is particularly significant, as most previously studied glutenases are evaluated solely for their activity against the 33-mer, with little attention given to their ability to process the resulting breakdown products nor does previous research provide a rationale for selection of the enzymes detailed therein^48–51^. Our data suggest that combining enzymes with complementary cleavage preferences can enable a more thorough and effective degradation of gluten peptides in the gastrointestinal tract. Such a strategy could prevent the accumulation of partially digested, still-immunogenic fragments, which represents a common limitation of single-enzyme therapies. Both enzymes are also active in the pH range of 4 to 6, thereby making them suitable for use in the full stomach/duodenum.

Together, these multi-parameter validations encompassing cytokine profiling, gene expression, barrier function, and confocal imaging underscore the biological efficacy of these enzymes in neutralizing gluten toxicity. Moreover, the complementary action of PSP464 on the 11-mer, a likely breakdown product of the 33-mer, supports a sequential degradation model wherein a cocktail of enzymes provides more complete detoxification than monotherapy. This layered approach not only reduces the concentration of intact immunogenic peptides but also limits the accumulation of shorter, still-immunogenic fragments, which are often overlooked in enzyme evaluations.

Gluten is a heterogeneous and structurally complex protein composite found primarily in wheat, barley, and rye, comprising prolamins (gliadins and hordeins) and glutelins. Its immunopathogenicity in Celiac disease (CeD) stems from the abundance of proline-and glutamine-rich motifs that resist proteolysis by human gastrointestinal enzymes. Notably, several immunogenic peptides such as the 33-mer from α-gliadin^6^, 18-mer^52^, 13-mer^53^, 9-mer^54^ and the 11-mer^15^ have been shown to activate gluten-reactive CD4⁺ T cells in HLA-DQ2/DQ8 individuals. Because of this structural and immunological complexity, a single enzyme may be insufficient to achieve complete detoxification. Our findings demonstrate that a data-driven, structure-guided strategy can successfully identify complementary enzymes such as PSP692 and PSP464 that act on different immunogenic peptide substrates. Although these enzymes show promising activity against model gliadin epitopes in vitro, further studies are warranted to evaluate their utility in gluten degradation and potential to reduce antigenic burden in the context of CeD management as a preventive or adjunctive strategy to reduce inadvertent exposure.

Beyond the two enzymes characterized in this study, the human gut microbiome harbors a far greater diversity of uncharacterized peptidases with potential gluten-degrading activity. Given the complexity of gluten immunopathogenicity, which involves multiple overlapping epitopes and resistant peptide fragments, systematic mining of metagenomic data could yield additional candidates with complementary substrate specificities, broader pH stability, or enhanced catalytic efficiency. However, a key limitation of the present study is the reliance on the limited number of publicly available CeD-associated metagenomes, which constrains the diversity of enzymes that can be identified. The paucity of high-quality, well-annotated metagenomic datasets from CeD and non-CeD cohorts remains a major bottleneck for comprehensive enzyme discovery. Expanding global efforts to generate large, clinically stratified metagenomic resources will therefore be critical for uncovering the full enzymatic repertoire of the gut microbiome and for developing next-generation combinatorial strategies to mitigate gluten immunotoxicity in vivo.

## Conclusion

The approach described herein integrating metagenomic mining, structural modeling, docking, and functional assays offers a robust pipeline for identifying additional gluten-degrading enzymes from human gut microbial communities. Given the diverse array of gluten epitopes and the variability in individual microbiomes, extending this approach to broader datasets may uncover novel enzyme classes with improved specificity, pH stability, or proteolytic efficiency. Future research should aim to establish the in vivo efficacy of such enzymes in more complex physiological contexts, including gastrointestinal transit, exposure to host proteases, and interactions with the host immune system. Rather than focusing exclusively on recombinant enzyme supplementation, synthetic biology-based strategies could also be explored. For instance, stable introduction of validated glutenase genes into commensal duodenal bacteria (e.g., *Prevotella spp*, *Faecalibacterium prausnitzii*, or *Lactobacillus spp*.) could enable sustained, host-compatible gluten detoxification at the site of exposure^55^. This would align with the long-term goal of a holistic microbiome-mediated management of gluten immunotoxicity in CeD.

## Abbreviations

Ni-NTA: Ni^2+^-nitrilotriacetate
DPP: dipeptidyl peptidase
p-NA: para-nitroanilide
Z: benzyloxycarbonyl

## Acknowledgements

The authors would like to acknowledge Department of Biotechnology (DBT), Govt. of India and National Centre for Cell Science for providing all necessary facilities to carry out research. BK was supported by a fellowship from DBT for the tenure of this research work.

## Data availability Statement

All metagenomic datasets analyzed in this study were obtained from publicly available repositories. Specifically, we accessed raw sequencing data from NCBI BioProject IDs PRJNA757365 and PRJNA486782. Supplementary sequence files and protein models generated during this study are included in the supplementary materials of the manuscript. No new sequencing data were generated. Additional processed files, including contigs encoding putative S8/S9/Xaa-Pro domain proteins, are provided in Supplementary Table ST1. All other supporting data are available upon reasonable request from the corresponding author.

